# Genome-wide association study of cryptosporidiosis in infants implicates *PRKCA*

**DOI:** 10.1101/819052

**Authors:** Genevieve L. Wojcik, Poonum Korpe, Chelsea Marie, Josyf Mychaleckyj, Beth D. Kirkpatrick, Stephen S. Rich, Patrick Concannon, A. S. G. Faruque, Rashidul Haque, William A. Petri, Priya Duggal

**Author notes:** Address correspondence to Priya Duggal,.

## Abstract

Diarrhea is a major cause of both morbidity and mortality worldwide, especially among young children. Cryptosporidiosis is a leading cause of diarrhea in children, particularly in South Asia and Sub-Saharan Africa where it is responsible for over 200,000 deaths per year. Beyond the initial clinical presentation of diarrhea, it is associated with long term sequelae such as malnutrition and neurocognitive developmental deficits. Risk factors include poverty and overcrowding, yet not all children with these risk factors and exposure are infected, nor do all infected children develop symptomatic disease. One potential risk factor to explain these differences is their human genome. To identify genetic variants associated with symptomatic cryptosporidiosis, we conducted a genome-wide association study (GWAS) examining 6.5 million single nucleotide polymorphisms (SNPs) in 873 children from three independent cohorts in Dhaka, Bangladesh: the Dhaka Birth Cohort (DBC), the Performance of Rotavirus and Oral Polio Vaccines in Developing Countries (PROVIDE) study, and the Cryptosporidiosis Birth Cohort (CBC). Associations were estimated separately for each cohort under an additive model, adjusting for height-for-age Z-score at 12 months of age, the first two principal components to account for population substructure, and genotyping batch. The strongest meta-analytic association was with rs58296998 (P=3.73×10^−8^), an intronic SNP and eQTL of *PRKCA*. Each additional risk allele conferred 2.4 times the odds of cryptosporidiosis in the first year of life. This genetic association suggests a role for protein kinase C alpha in pediatric cryptosporidiosis and warrants further investigation. This article was submitted to an online preprint archive.(1)

**Importance:** Globally, one of the major causes of pediatric morbidity and mortality remains diarrhea. The initial symptoms of diarrhea can often lead to long term consequences for the health of young children, such as malnutrition and neurocognitive developmental deficits. Despite many children having similar exposures to infectious causes of diarrhea, not all develop symptomatic disease, indicating a possible role for human genetic variation. Here we conducted a genetic study of susceptibility to symptomatic disease associated with Cryptosporidium infection (a leading cause of diarrhea) in three independent cohorts of infants from Dhaka, Bangladesh. We identified a genetic variant within protein kinase c alpha (*PRKCA*) associated with higher risk of cryptosporidiosis in the first year of life. These results indicate a role for human genetics in susceptibility to cryptosporidiosis and warrant further research to elucidate the mechanism.

## Introduction

Cryptosporidiosis is a leading cause of diarrhea and is estimated to be responsible for greater than 200,000 deaths in young children in South Asia and Sub-Saharan Africa each year. (2) Beyond the immediate infection, cryptosporidiosis is also associated with long term sequelae including malnutrition and neurocognitive developmental deficits. (3–6) The majority of human infections are caused by the *C. hominis, C. meleagridis,* and *C. parvum* species, members of the phylum Apicomplexa. (5, 7, 8) As cryptosporidiosis is transmitted fecal-orally, contact with any reservoir with possible fecal contamination could serve as point of transmission. In the developed world, cryptosporidia are an important cause of diarrhea in individuals living with HIV and is the most common pathogen causing waterborne outbreaks. (8)

In endemic regions, cryptosporidiosis mostly impacts young children, and risk factors for infection include poverty, and overcrowding. (5, 9–11) Livestock serve as an environmental reservoir for *C. parvum*, and transmission has been reported after contact with infected animals or drinking water contaminated by human or animal waste. (12) In regions where *Cryptosporidium* infection is endemic, there is heterogeneity in clinical course and outcome. In an eight-site multicenter international study of enteric infection and malnutrition (MAL-ED), the rate of *Cryptosporidium* infection, age of onset, number of repeat infections, and clinical manifestation varied significantly by site. (10) In a recent study in Dhaka, Bangladesh, we found that two-thirds of children living in an urban slum were infected with *Cryptosporidium* by two years of age and one-fourth had more than one episode of cryptosporidiosis. Fully three-fourths of infections were subclinical, but regardless of symptoms, children with cryptosporidiosis were more likely to become malnourished by age two years. (5) Potential explanations for the *Cryptosporidium* infection heterogeneity include differences in pathogenicity of various *Cryptosporidium* species or genotypes (13), as well as host genetic susceptibility.

Candidate gene studies identified an increased risk of *Cryptosporidium* infection associated with specific alleles in HLA class I and II genes and SNPs in the mannose binding lectin (*MBL*) gene. (14–16) Bangladeshi preschool children with multiple *Cryptosporidium* infections (≥2 infections) were more likely to carry the -221 *MBL2* promoter variant (rs7906206; OR=4.02, P=0.025) and have the YO/XA haplotype (OR=4.91), as well as be deficient in their MBL serum levels (OR=10.45). (15) Since *MBL* and HLA alleles only partially explained *Cryptosporidium* susceptibility, we conducted a genome-wide association study (GWAS) of cryptosporidiosis occurring in the first year of life using three existing birth cohorts of children in Dhaka, Bangladesh: the Performance of Rotavirus and Oral Polio Vaccines in Developing Countries (PROVIDE) study, the Dhaka Birth Cohort (DBC), and the Cryptosporidiosis Birth Cohort (CBC).

## Results

Across these three cohorts, there were a total of 183 children with at least one symptomatic (diarrheal) sample that tested positive for *Cryptosporidium* within the first year of life (“cases”). (**Table 1**) A total of 873 children did not test positive for *Cryptosporidium* in either symptomatic (diarrheal) or surveillance samples within the first year of life (“controls”). There were no significant differences in height-for-age Z-score (HAZ) at birth (HAZ_birth_), the number of days exclusively breastfed, or sex between cases and controls (P>0.05). To control for a possible role of malnutrition affecting susceptibility to infection, we compared the HAZ at 12 months of age (HAZ_12_) between cases and controls. While we observed increased levels of stunting (lower HAZ_12_) within PROVIDE (*P*=0.007), we observed no difference with the other two cohorts (P>0.05). Additionally, there was no statistically significant evidence of heterogeneity in HAZ_12_, number of days exclusively breastfed, or sex between the three studies (P_het_>0.05).

**Table 1:** Demographics of study populations.

| | Dhaka Birth Cohort (DBC) | | | PROVIDE | | | Crypto Cohort | | | $P_{het}$ |
| --- | --- | --- | --- | --- | --- | --- | --- | --- | --- | --- |
|  | Controls<br>(N=267)<br>Mean | Cases<br>(N=46)<br>Mean | P-value | Controls<br>(N=354)<br>Mean | Cases<br>(N=60)<br>Mean | P-value | Controls<br>(N=252)<br>Mean | Cases<br>(N=77)<br>Mean | P-value |  |
| **HAZ @ 12 months** | -1.75 | -1.74 | 0.97 | -1.40 | -1.79 | $7.28 \times 10^{-3}$ | -1.34 | -1.63 | 0.02 | 0.12 |
| **Exclusive Breastfeeding (days)** | 130.2 | 114.6 | 0.16 | 127.2 | 112.1 | 0.06 | 110.9 | 103.7 | 0.42 | 0.74 |
| **Sex (% Female)** | 46.3% | 34.8% | 0.15 | 45.9% | 46.7% | 0.91 | 52.8% | 57.7% | 0.45 | 0.28 |

### GWAS of *Cryptosporidiosis* within the first year of life

We tested the association between 6.5 million SNPs across the human genome with symptomatic *Cryptosporidium* infection in the first year of life. Effects were estimated separately for the three birth cohorts and subsequently combined using a fixed-effects meta-analysis, filtered for heterogeneity (P_het_), minor allele frequency (MAF) >5%, and imputation quality (INFO>0.6). (**Figure 1, Supp. Figure 1**) A total of 6 SNPs in an intron of *PRKCA* (protein kinase c, alpha) were significantly associated with *Cryptosporidium* infection (P<5×10^−8^). (**Figure 2A**) For the SNP most associated with *Cryptosporidium* infection (rs58296998), each copy of the risk allele (T) conferred 2.4 times the odds of cryptosporidiosis within the first year of life (P=3.73×10^−8^). This effect size and risk allele were consistent across all three studies (P_het_=0.11). (**Figure 2B**) After conditioning on rs58296998 (by including this SNP in the logistic regression model as a covariate), the evidence for association with the remaining SNPs in the region was no longer significant, suggesting that the observed association in *PRKCA* is explained by a single SNP (rs58296998) or one highly correlated with this SNP. (**Figure 2C**) Of the 26 children homozygous for the risk allele (TT) at rs58296998, 46% developed symptomatic *Cryptosporidium* during the first year of life. This proportion decreased to 24% for children heterozygous (CT) for this risk allele (N=272), compared to 13% of children homozygous (CC) for the non-risk allele (N=745).

**Figure 1:**
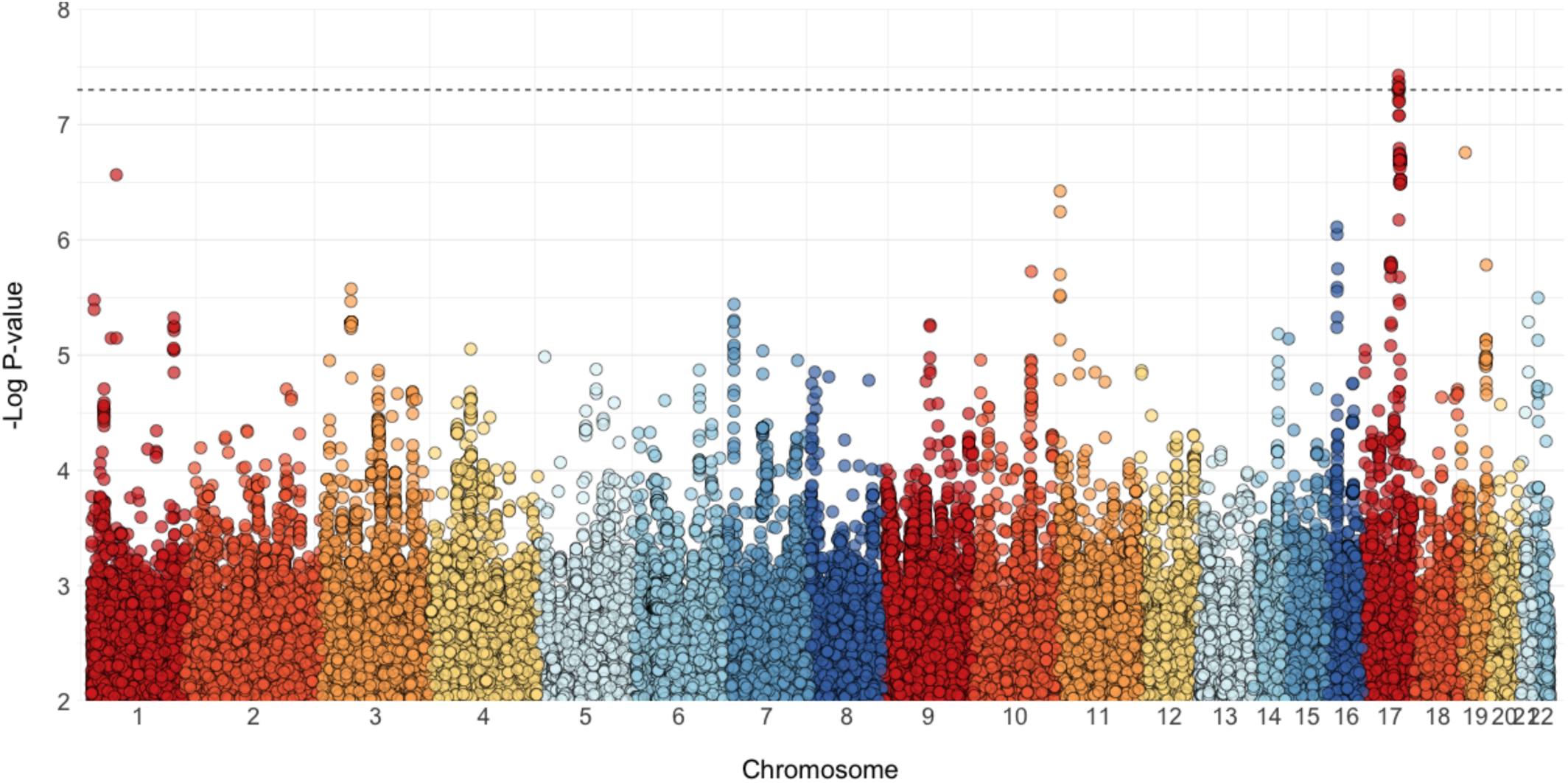
Manhattan Plot for Cryptosporidiosis within the first year of life. Each dot indicates the association of a single SNP with Cryptosporidiosis in the first year of life. SNPs are sorted by chromosome (each color) and position along the x-axis. The y-axis is the -log10 P-value for the SNP association the meta-analysis of study-specific logistic regressions adjusting for height-for-age Z-score at 12 months, the first two study-specific principal components, and batch for the Dhaka Birth Cohort (DBC). Genome-wide significance (5×10^−8^) is denoted by the dashed line. This plot is limited to associations with a P-value below 0.01.

**Figure 2:**
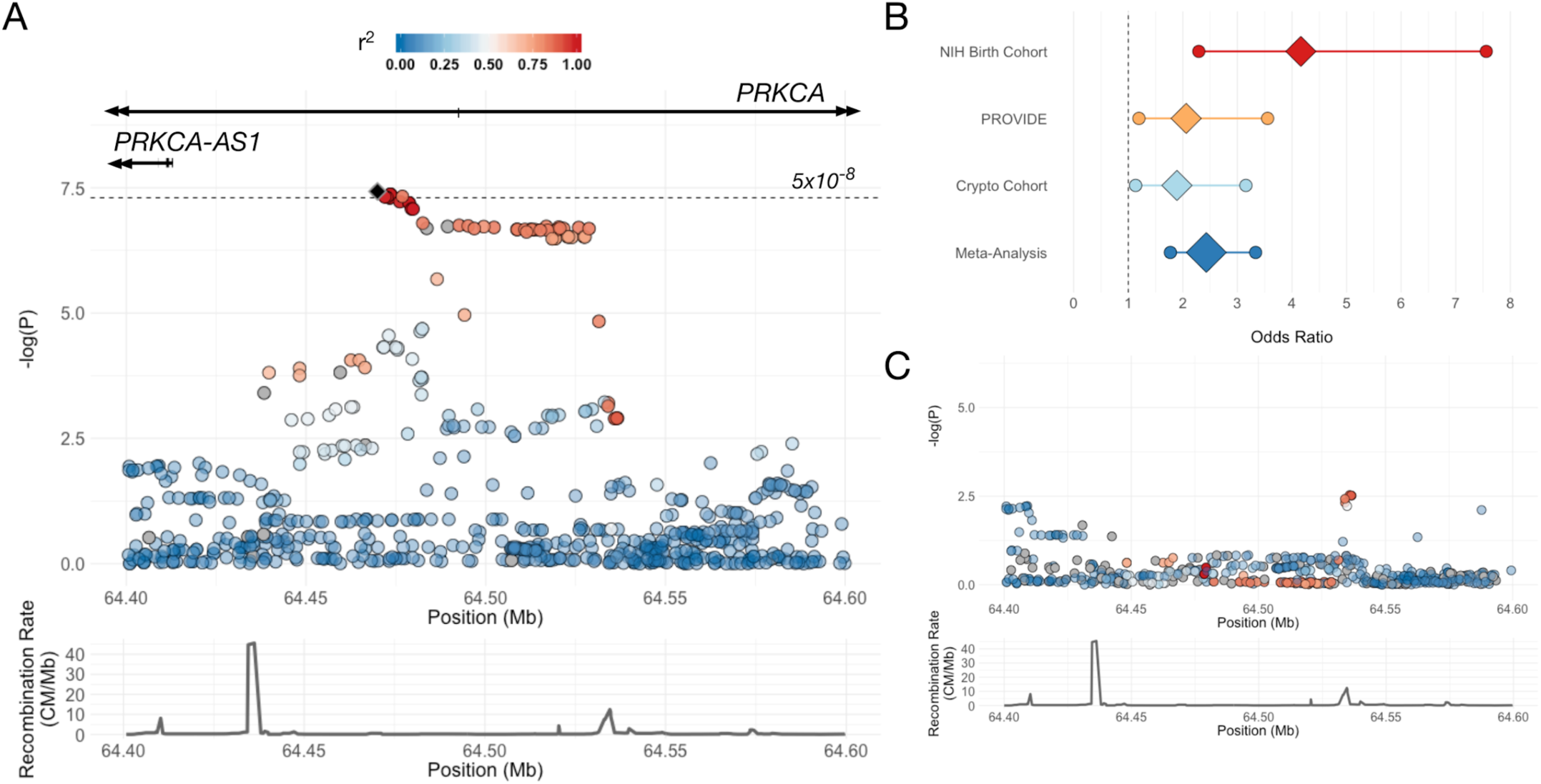
Association between variants in *PRKCA* and cryptosporidiosis. (A) Regional association on chromosome 17 between variants in *PRKCA* and cryptosporidiosis. Fill denotes linkage disequilibrium (r^2^) between the top SNP (rs58296998) and surrounding SNPs. (B) Forest plot of odds ratios and 95% confidence intervals for top signal rs58296998 by individual cohort and meta-analysis. (C) Regional association in PRKCA region after conditioning on top signal rs58296998, showing significantly diminished signal between recombination peaks.

The rs58296998 T allele frequencies for all three cohorts (15.0-16.7%) in this region are consistent with the Bangladeshi reference population (1000 Genomes Phase 3) of 18% and the overall South Asian frequency of 15%. (Auton et al. 2015) Globally, the highest frequencies of rs58296998 T allele are found in East Asian populations, with the highest T allele frequency of 34% of the Chinese Dai in Xishuangbanna, China. The rs58296998 T allele is at lower frequencies within Africa, at 9% within the Luhya in Kenya, and even less frequent in West Africa (3.5-5.5%). (**Figure 3**)

**Figure 3:**
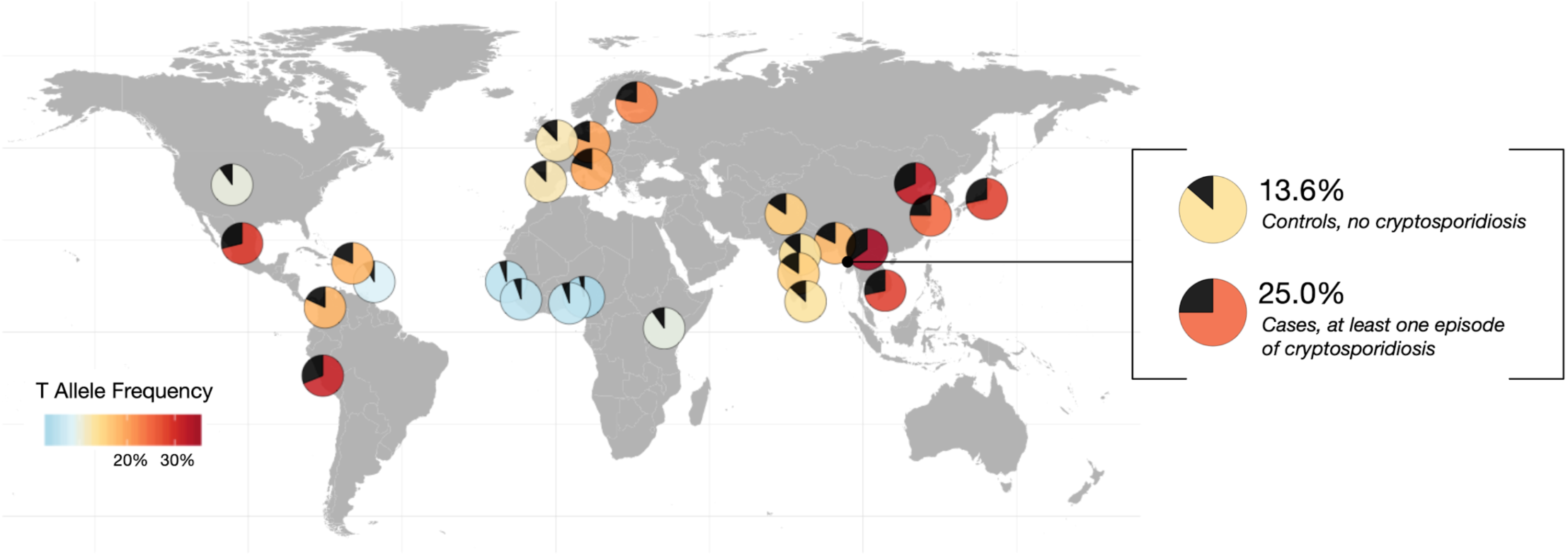
Allele frequencies for allele T at top signal rs58296998 from 1000 Genomes Phase 3 data, as well as by case/control status in three cohorts combined. Each pie chart on the map shows the frequency of the T allele with the black wedge. The remainder of the pie chart is colored by that T allele frequency. The inset provides the T allele frequency for children without any symptomatic cryptosporidiosis in the first year of life (controls; MAF=13.6%) and those with at least one diarrheal episode (cases; MAF=25.0%).

Cases had their first diarrheal episode positive for *Cryptosporidia* at a mean of 242 days of age. We confirm the GWAS results, with the dosage of rs58296998 risk alleles significantly associated with time to first diarrheal sample positive for *Cryptosporidia* among cases versus right-censored controls (up to the child’s first birthday) (P=6.37×10^−8^). All children homozygous for the risk allele (TT) have their first episode in the first year of life. (**Supp. Figure 2**) Among cases, however, we there was no statistically significant association between rs58296998 genotype and time to infection (P=0.095). In PROVIDE, the rs58296998 genotype was associated with severity of diarrhea as determined by the Ruuska score (*P*=0.028). (**Supp. Figure 3**)

Suggestive SNP associations with *Cryptosporidium* (*P*<10^−6^) were also identified on chromosomes 11 and 16. The strongest association on chromosome 11 (rs4758351) is located within an intergenic region of a cluster of olfactory receptor genes. Each copy of the rs4758351 A allele (MAF:14%) conferred 2.39 times the odds of *Cryptosporidium* within the first year of life (P=3.78×10^−7^). **(Supp. Figure 4)** Multiple SNPs in this region of chromosome 11 (chr11:6,015,194-6,024,551) had similar magnitude and strength of association with *Cryptosporidium* (OR: 2.13-2.39). The strongest association on chromosome 16 was with the rs9937140 SNP, located upstream of apolipoprotein O pseudogene 5 (*APOOP5*). Each copy of the rs9937140 G allele (MAF: 23%) conferred 1.99 times the odds of cryptosporidiosis (P=7.75×10^−7^). **(Supp. Figure 5)**

**Figure 4:**
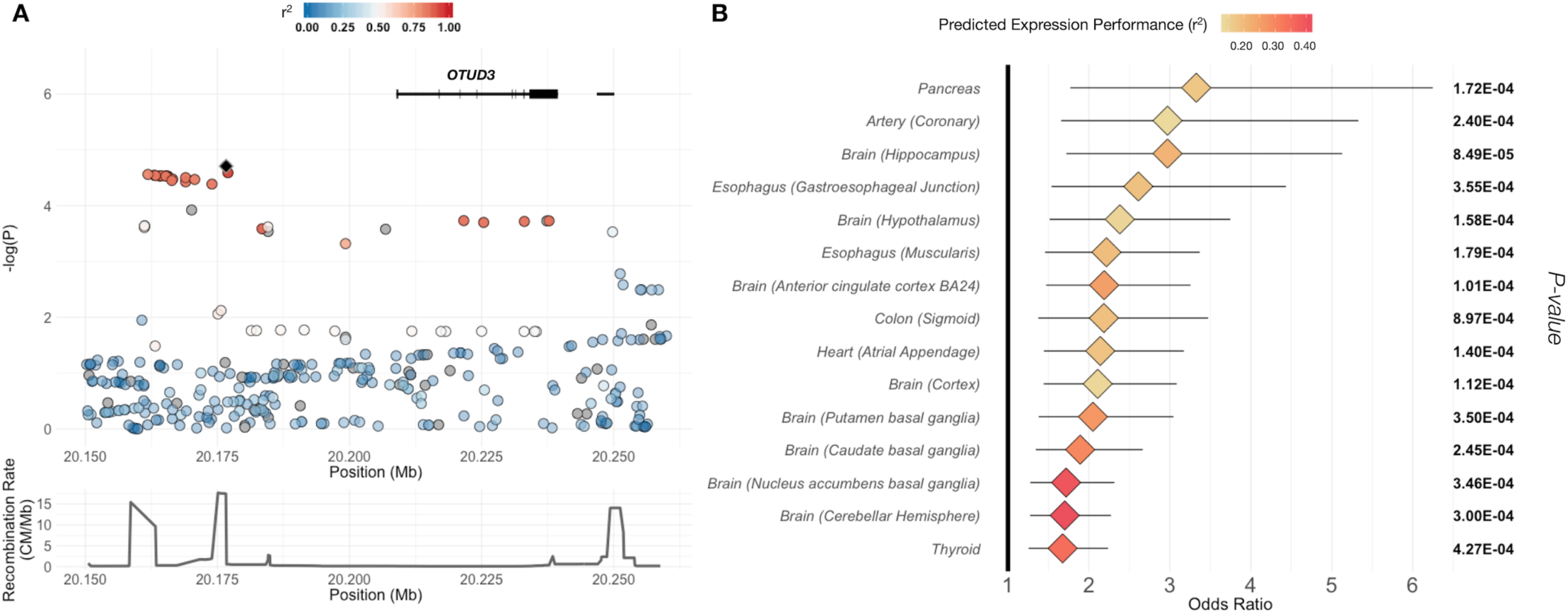
*OTUD3* region showing association with cryptosporidiosis in the first year of life. (A) Association of SNPs on chromosome 1 region, colored by linkage disequilibrium (r^2^) with index SNP (black diamond). (B) Association of case status with imputed gene expression in all tissues with P<0.001 and predicted expression performance of r^2^>0.1.

### Expression and PrediXcan

We used the publicly available resource, Genotype-Tissue Expression (GTEx) project to estimate the influence of human genetic variation on human gene expression in multiple tissues. (17, 18) The associated rs58296998 SNP, located in the *PRKCA* gene, is also associated with *PRKCA* expression. This expression quantitative trait locus, eQTL, showed decreasing expression of *PRKCA* with each T allele in esophageal muscularis (*P*= 3.12×10^−5^), the sigmoid colon (*P*=4.61×10^−4^), and esophageal mucosa (*P*=7.50×10^−4^). (18) These expression data, coupled with the GWAS result, suggests that decreased expression of *PRKCA* is correlated with increased risk of symptomatic *Cryptosporidium* infection within the first year of life.

### Additional Genome-Wide Expression and Gene Set Analyses

In the absence of direct gene expression measurement, we relied on previously estimated tissue-specific associations between genome-wide SNPs and gene expression, which quantify the genetic component of gene expression. We estimated predicted patterns of genome-wide differential gene expression between cases and controls by weighting the summary statistics from our GWAS of cryptosporidiosis in the first year of life by tissue-specific PredictDB weights. These SNP-level estimates were then combined for each gene to infer association between imputed gene expression and *Cryptosporidiosis*. (19)(20) No association of predicted gene expression with cryptosporidiosis reached statistical significance. A total of 13 genes had nominally significant (P< 0.001) association in more than one tissue-specific model. **(Supp. Table 1, Supp. Figure 6)** Variants in the gene *OTUD3* (OTU deubiquitinase 3, chr1:20,208356-20,239,438) were associated with cryptosporidiosis in 18 different tissue-specific models at P<0.001. In all tissue-specific models, individuals with predicted increased expression of *OTUD3* had an increased risk of cryptosporidiosis within the first year of life (OR: 1.68-6.63, P: 8.46×10^−5^ - 8.97×10^−4^). **(Figure 4)**

We also performed gene set enrichment analysis using MSigDB hallmark gene sets (N=50), KEGG (N=186) and BioCarta (N=217) by combining gene-level summary statistics to examine aggregate signals within biological pathways. No pathways reached statistical significance after adjusting for multiple comparisons; however, several gene sets were suggestive. (**Supp. Table 2**) The two top-ranked gene sets are part of the hedgehog signaling pathway: the hallmark hedgehog signaling (P_emp_=5.04×10^4^, BF=515.65) and KEGG hedgehog signaling pathway (P_emp_=1.47×10^−3^, BF=235.59).

## Discussion

Here we present the results of the first genome-wide association study of symptomatic *Cryptosporidium* infection. Specifically, we tested the role of host genetics in the susceptibility to *Cryptosporidium* infection associated with diarrhea within the first year of life. A region on chromosome 17 was identified, with each additional T allele of rs58296998, an intronic SNP in *PRKCA*, conferring 2.4 times the odds of cryptosporidiosis within the first year of life. Additionally, this SNP was previously identified as an eQTL of *PRKCA*, with decreased expression of *PRKCA* associated with the T allele. This suggests that this SNP may influence *Cryptosporidium* infection through decreased expression of *PRKCA*.

*Protein kinase c alpha (PRKCA)* is an isotype of the protein kinase C (PKC) family, which are serine- and threonine-specific and known to be involved in diverse cellular signaling pathways. Specifically, PKCs have numerous roles in the development and function of the gastrointestinal tract (21) and in the immune response (22). This relationship was confirmed with knockout experiments, where PKCα was shown to be a positive regulator of Th17 cell effector functions. PKCα-deficient (Prkca(-/-)) cells failed to mount the appropriate levels of IL-17A i*n vitro*. (22) An analysis of *Cryptosporidium parvum-*infected mice demonstrated the importance of the Th17 response to infection, showing increased levels of IL-17 mRNA and Th17 cell-related cytokines in gut tissue after infection. (23) Additionally, both pharmacological and genetic PKCα inhibition have been shown to prevent NHE3 internalization, Na+ malabsorption, and TNF-mediated diarrhea, despite continued barrier dysfunction (24), supporting a role for *PRKCA* in symptomatic cryptosporidiosis. This link between *PRKCA* and Th17 may be critical to gut infections, and specifically to infection of Cryptosporidium in the developing infant gut. We identified a SNP associated with decreased expression of *PRKCA* and thus less able to mediate the IL-17 immune response during Cryptosporidium infection. *PRKCA* has also been shown to be associated with numerous other infections, including *Staphylococcus aureus* (25), progression of sepsis (26), toxoplasmosis (27), *Burkholderia cenocepacia* in cystic fibrosis patients (28), and hepatitis E virus replication (29).

As an obligate intracellular parasite, *Cryptosporidium* relies on host cells to complete its life cycle in the human host, thus it is also plausible that *PRKCA* may directly mediate susceptibility via impacts on parasite invasion. Sporozoites invade brush border intestinal epithelial cells by inducing volume increases (30) and cytoskeletal remodeling at the site of host cell attachment (31) which leads to engulfment via host membrane protrusions. Studies have shown that inhibition of host factors, including actin remodeling proteins and PKC enzymes, is sufficient to inhibit sporozoite invasion *in vitro*. (31) Interestingly, PKCα has been shown to play an important role in *Escherichia coli* pathogenesis.(32) Like *Cryptosporidium, E. coli* induces host actin condensation at the site of host cell invasion and immunocytochemical studies indicate that activated PKCα co-localized with actin condensation at the bacterial entry site.(33)

While our top SNP within *PRKCA* has previously been shown to influence the expression of *PRKCA* in GTEx, our imputed gene expression analysis using PrediXcan did not see a significant difference in predicted *PRKCA* expression between cases and controls. This is likely due to the difference between a single SNP being examined in GTEx versus the combined effects of multiple eQTLs estimated from a European descent reference population in PrediXcan. A major limitation of predicted gene expression analyses is the lack of population-specificity for non-European groups. (34) The PrediXcan models were derived from European-descent individuals, as were the covariance structures used to infer correlation between eQTLs. We see a direct relationship between population differences in allele frequencies for the weighted SNPs and impaired performance. Specifically, we observe the lowest predictive performance in tissues for which the informative SNPs have large differences in allele frequencies between European and South Asian populations in the 1000 Genomes Project phase 3 data (35). (**Supp. Figure 7**) These include two tissues, esophageal mucosa and the colon sigmoid tissue, in which rs58296998 was identified as an eQTL for *PRKCA*. These trends highlight the importance of reference populations representative of global populations to ensure tools are useful in non-European populations, such as ours. We also identified increased expression of *OTUD3* to be associated with increased odds of cryptosporidiosis within the first year of life. This gene is associated with ulcerative colitis (36–42) and inflammatory bowel disease (43, 44). This finding is consistent with a shared pathway between enteric infection and autoimmune intestinal disease, as indicated in a previous genetic analysis of *Entamoeba histolytica* infection in the same study population. (45)

When the predicted patterns of differentially expressed genes are collapsed into gene sets, we found enrichment in the hedgehog signaling pathway. A previous study examined the gene expression profiles of long non-coding RNA (lncRNA) and mRNA in HCT-8 cells infected with *C. parvum* IId subtype. (46) Of note, *PRKCA* was the most significantly differentially expressed gene in infected HCT8 cells 24 hours post infection (2.24-fold decreased expression in infected cells P=3.82×10^−5^). Pathway analysis of the differentially expressed mRNAs found that genes in the hedgehog signaling pathway were significantly enriched during *Cryptosporidium* infection. This finding in combination with our identification of hedgehog signaling in imputed gene expression profiles is suggestive of a potential link between decreased *PRKCA* expression and hedgehog signaling, however further research to confirm these findings and elucidate the role of genetic variation in *PRKCA* on gene expression and hedgehog pathway perturbation is needed.

A potential limitation of our study is that due to the use of sensitive molecular diagnostics multiple enteropathogens were frequently detected in each diarrheal sample. However, we did not detect the same genetic signatures as from our previous study of *Entamoeba histolytica* in this same study population for Cryptosporidium. (45) Therefore, we are confident that our results are specific to cryptosporidiosis, despite co-occurrence with other enteric pathogens.

Through a GWAS meta-analysis of three separate birth cohorts, we identify a region in *PRKCA* on chromosome 17 as being associated with increased risk of symptomatic cryptosporidiosis in the first year of life among Bangladeshi infants. This gene has previously been implicated in other infectious outcomes, indicating pleiotropy with the immune system’s reaction to numerous pathogens. Publicly available data supports a link between our top SNP and expression of *PRKCA*, suggesting a mechanism via Th17 inflammatory control. Clinical trials are currently be proposed for PKC isotypes, including PKC-alpha, for autoimmune disease, and therefore may be important for cryptosporidiosis which lacks treatment for young children. (47) Identifying host genetic variation associated with cryptosporidiosis, like PRKCA, can help us identify viable drug targets to improve treatment and prevention of this major cause of morbidity and mortality. Further research is needed to elucidate the mechanism underlying this relationship and to better understand the complex interplay of genetic susceptibility and environmental influences in the development of intestinal disease.

## Methods

The study protocol was approved by the Research and Ethical Review Committees of the International Center for Diarrheal Disease Research, Bangladesh, and the Institutional Review Boards of the University of Virginia and the Johns Hopkins Bloomberg School of Public Health. The parents or guardians of all individuals provided informed consent.

### Dhaka Birth Cohort study design

Designed to study the influence of malnutrition in child development, the Dhaka Birth Cohort (DBC) is a subset of a larger birth cohort recruited from the urban slum in the Mirpur Thana in Dhaka, Bangladesh. Children were enrolled within the first week after birth and followed-up bi-weekly with household visits by trained field research assistants for the first year of life. Anthropometric measurements were collected at the time of enrollment and every three months thereafter. Height-for-age adjusted Z-scores (HAZ) scores were calculated by comparing the height and weight of study participants with the World Health Organization (WHO) reference population, adjusting for age and sex, with WHO Anthro software, version 3.0.1. Field research assistants (FRAs) collected diarrheal stool samples from the home or study field clinic every time the mother of the child reported diarrhea. To maintain a cold chain, the samples were transported to the Centre for Diarrhoeal Disease Research, Bangladesh (ICDDR,B) parasitology laboratory. The presence of Cryptosporidium was determined using enzyme-linked immunosorbent assay (ELISA). More details can be found in Steiner et al (2018) and Korpe et al (2018). (5, 10) We used a nested case-control design, where children with at least one diarrheal sample positive for Cryptosporidium within the first year were defined as “cases”. Children with diarrheal samples, of which none are positive for *Cryptosporidium*, were defined as “controls”.

### PROVIDE study design

The “Performance of Rotavirus and Oral Polio Vaccines in Developing Countries” (PROVIDE) Study is a randomized controlled clinical trial and birth cohort also from the same urban slum in the Mirpur Thana in Dhaka, Bangladesh as the DBC and Cryptosporidia Cohort (below). PROVIDE was specifically designed to assess the influence of various factors on oral vaccine efficacy among children in areas with high poverty, urban overcrowding, and poor sanitation. The 2 ×2 factorial design looked specifically at the efficacy of the 2-dose Rotarix oral rotavirus vaccine and oral polio vaccine (OPV) with an inactivated polio vaccine (IPV) boost over the first two years of life. All participants were from the Mirpur area of Dhaka, Bangladesh, with pregnant mothers recruited from the community by female Bangladeshi FRAs. Each participant had fifteen scheduled follow-up clinic visits, as well as biweekly diarrhea surveillance through home visits by FRAs. The presence of Cryptosporidium in diarrheal samples was determined by ELISA. Consistently with the DBC phenotype definition, cases had at least one diarrheal sample positive for Cryptosporidium within the first year of life. Controls had at least one diarrheal sample available for testing, but none were positive for Cryptosporidium. Severity of diarrheal was determined with the Ruuska score, which assesses severity as a function of diarrhea length, clinical symptoms, and other clinical features. (48)

### Cryptosporidia Cohort study design

The Cryptosporidia cohort (“Cryptosporidiosis and Enteropathogens in Bangladesh”; ClinicalTrials.gov identifier NCT02764918) is a prospective longitudinal birth cohort study in two sites in Bangladesh. The first is in a urban, economically depressed neighborhood of Mirpur, and the second is in a rural subdistrict 60 km northwest of Dhaka, called Mirzapur. The two birth cohorts were established in parallel, with the objective of understanding the incidence of cryptosporidiosis, the acquired immune response, and host genetic susceptibility to cryptosporidiosis in Bangladeshi children. Pregnant women were recruited and screened, and infants were enrolled at birth. Participants were followed twice-weekly with in-home visits to monitor for child morbidity and diarrhea for 24 months. Infant height and weight were measured every 3 months, and weight-for-age and height-for-age adjusted z-scores were determined using World Health Organization Anthro software (version 3.2.2). Stool samples were collected during diarrheal illness and once per month for surveillance. Stool was tested for Cryptosporidium by quantitative polymerase chain reaction (qPCR) assay modified from Liu et al. (49) A cycle threshold of 40 was used. The pan-Cryptosporidium primers and probes target the 18S gene in multiple species known to infect humans. (5)

### Genotype data

The Dhaka Birth Cohort (DBC) and PROVIDE Study data was generated and cleaned as described previously. (45) A summary of quality control procedures are detailed in **Supp. Figure 1**. Briefly, a total of 396 children in DBC were genotyped on three different Illumina arrays. All individuals were imputed to 1000Genomes Phase 3 data. After post-imputation QC, which included additional filtering for relatedness and poorly imputed variants, a total of 396 individuals and 10.2 million SNPs were included in the DBC data freeze. For PROVIDE, a total of 541 individuals were genotyped on Illumina’s Multi-ethnic Genotyping Array (MEGA). After standard quality control measures, including minor allele frequency >0.5%, missingness <5%, and first degree related individuals removed, a total of 499 individuals remained. After imputation to 1000Genomes and subsequent post-imputation QC, a total of 499 individuals and 10.8 million genetic variants remained. For the Cryptosporidium Cohort, a total of 630 individuals were genotyped on Illumina’s Multi-ethnic Global Array (MEGA). One individual was removed for first-degree relatedness (PI_HAT>0.2), 31 individuals removed as PCA outliers, and 3 individuals were removed for heterozygosity. No individuals or SNPs were removed for missingness (>5%). Additional SNP-level filters included minor allele frequency (MAF)<0.5% (M=751,869) and Hardy-Weinberg equilibrium P-value<10^−5^ (M=85). After all QC steps, CryptoCohort genotype data included 594 individuals and 826,228 SNPs. Phasing in SHAPEIT2 (50) was followed by imputation with IMPUTE2 (51, 52) to 1000 Genomes Phase 3 data (1000Genomes). (35) All three studies were separately imputed to 1000Genomes.

### Cross-study genetic data harmonization

After imputation, all three datasets (DBC, PROVIDE, CryptoCohort) were double checked for relatedness both within study, as well as between studies, to ensure independence. One individual from each pair of relateds were dropped consistent with up to second degree of relatedness (PI_HAT>0.2). Individual outliers for heterozygosity (F > 5 standard deviations from mean) were also excluded from further analysis. A total of 85 individuals were dropped from DBC, 9 from PROVIDE, and 34 from CryptoCohort. Only the top principal component from the combined dataset was found to be significantly associated with outcome. (**Supp. Figures 8-10**)

### Statistical analysis

Each study (DBC, PROVIDE, and CryptoCohort) was analyzed separately using logistic regression with an additive model accounting for imputed genotype weights in SNPTEST (51, 53, 54). All three analyses were adjusted for height-for-age Z-score (HAZ) at one year of age, sex, and the first two principal components. The Dhaka Birth Cohort was additionally conditioned on genotyping array to account for batch effects. We combined the three analyses in a fixed-effects meta-analysis within META. Results were filtered for P_het_>0.05, minor allele frequency (MAF) >5%, and INFO>0.6 in all three studies, resulting in 6,504,706 SNPs. The conditional analyses were run separately by cohort for the *PRKCA* region, each analysis conditioning on rs58296998 in addition to the original covariates with SNPTEST. Results were again filtered for heterogeneity or P(het)>0.05, MAF>5%, and INFO>0.6 in all three studies.

### Allele frequencies

The allele frequencies were derived from the 1000 Genomes Project Phase 3 data, v5a. (35) Individuals were stratified by their denoted population with first degree related individuals removed.

### GTEx and eQTL overlap with GWAS results

Expression quantitative trait loci (eQTLs) were identified through the GTEx Portal (https://www.gtexportal.org/home/) on August 6^th^, 2018. (18) The top SNP was identified as an eQTL for *PRKCA* with *P*<0.001 for multiple tissues.

### MetaXcan imputation and association analysis

To impute gene expression and association with outcome from our GWAS summary statistics, we applied MetaXcan (S-PrediXcan and packaged best practices). (20) Weights were previously derived with GTEx v7 data in a European-descent population, with accompanying European-descent linkage disequilibrium metrics for the SNP covariance matrices (PredictDB Data Respository: http://predictdb.org/). MetaXcan was used instead of the original PrediXcan to ensure consistency in models with our GWAS. All 48 tissues were run separately over the meta-analysis results previously described. Following imputation and estimation of gene expression with outcome, we calculated weights for each gene-tissue pair as the ratio between the number of SNPs used in the model versus the total number that were pre-specific in the model, multiplied by predicted expression performance. To determine associations across many tissues, a P-value threshold of 0.001 was utilized. A strict Bonferonni correction for the 242,686 comparisons results in a P-value threshold of 0.05/242,686=2.06×10^−7^, which no comparison yielded a statistically significant result. The relationship of allele frequencies in European and South Asian populations with PrediXcan weights were examined to assess prediction capacity. (**Supp. Figures 6, 11**)

### Gene set enrichment analysis

Gene set enrichment analysis was conducted on the imputed gene expression data summary statistics previously described from MetaXcan. For each gene, we selected the tissue with the smallest P-value. Using the program GIGSEA (Genotype Imputed Gene Set Enrichment Analysis (55)), we tested for association of 453 curated gene sets defined by MSigDB hallmark gene sets (56), as well as KEGG (Kyoto Encyclopedia of Genes and Genomes; www.kegg.jp) and BioCarta (57) gene sets. (58) To account for redundancy with overlapping gene sets, we utilized the weighted multiple linear regression model, using the matrix operation to increase speed, with a 1,000 permutations. A false discovery rate of 0.05 was calculated on the ranked results.

### Data and Code Availability

Data is publicly available from the NIH, via dbGAP, phs001478.v1.p1 Exploration of the Biologic Basis for Underperformance of Oral Polio and Rotavirus Vaccines in Bangladesh or by request from the authors. All analysis programs used are detailed above, but the actual code in R for each analysis is also available by request from the authors.

## Acknowledgements

The Genotype-Tissue Expression (GTEx) Project was supported by the <u>Common Fund</u> of the Office of the Director of the National Institutes of Health, and by NCI, NHGRI, NHLBI, NIDA, NIMH, and NINDS. This work was funded by grants to WP from the Bill & Melinda Gates Foundation and the National Institutes of Health, Allergy and Infectious Disease AI043596 and the Henske Family, and to PD from the Sherrilyn and Ken Fisher Center for Environmental Infectious Diseases Discovery Program. The funders had no role in the study design, data collection and data analysis, decision to publish, or preparation of the manuscript. Icddr,b is grateful to the governments of Bangladesh, Canada, Sweden, and the UK for providing core unrestricted support. We thank the families of the Mirpur field area who participated in this study, and we also thank the world of the field and lab staffs of the Parasitology Laboratory of icddr,b who worked for the Dhaka Birth Cohort (DBC), PROVIDE, and Crypto Birth Cohort (CBC) projects, without whom we could not have completed this research.

