## Supplementary Figures for "Genome-wide association study of cryptosporidiosis in infants implicates *PRKCA*"

Supplementary Figure 1: Quality Control workflow for all three cohorts. 2

Supplementary Figure 2: Survival analysis of first episode of cryptosporidiosis by PRKCA rs58296998 genotype within the first year of life. 3

Supplementary Figure 3: Relationship between genotype for PRKCA SNP rs58296998 and severity of diarrhea as determined by Ruuska score within PROVIDE. 4

Supplementary Figure 4: LocusZoom of suggestive signal on chromosome 11 (rs4758351). 5

Supplementary Figure 5: LocusZoom of suggestive signal on chromosome 16 (rs9937140). 6

Supplementary Figure 6: Shared associations for predicted gene expression, filtered for gene-tissue pairs with P<0.001. 7

Supplementary Figure 7: Gene expression prediction characteristics of *PRKCA*. 8

Supplementary Figure 8: Distribution of three studies for principal components 1-5, colored by study. 9

Supplementary Figure 9: Distribution of three studies for principal components 1-5, colored by case status. 10

Supplementary Figure 10: Histogram of heterozygosity distribution by cohort. 11

Supplementary Figure 11: Gene expression prediction characteristics of *OTUD3*. 12

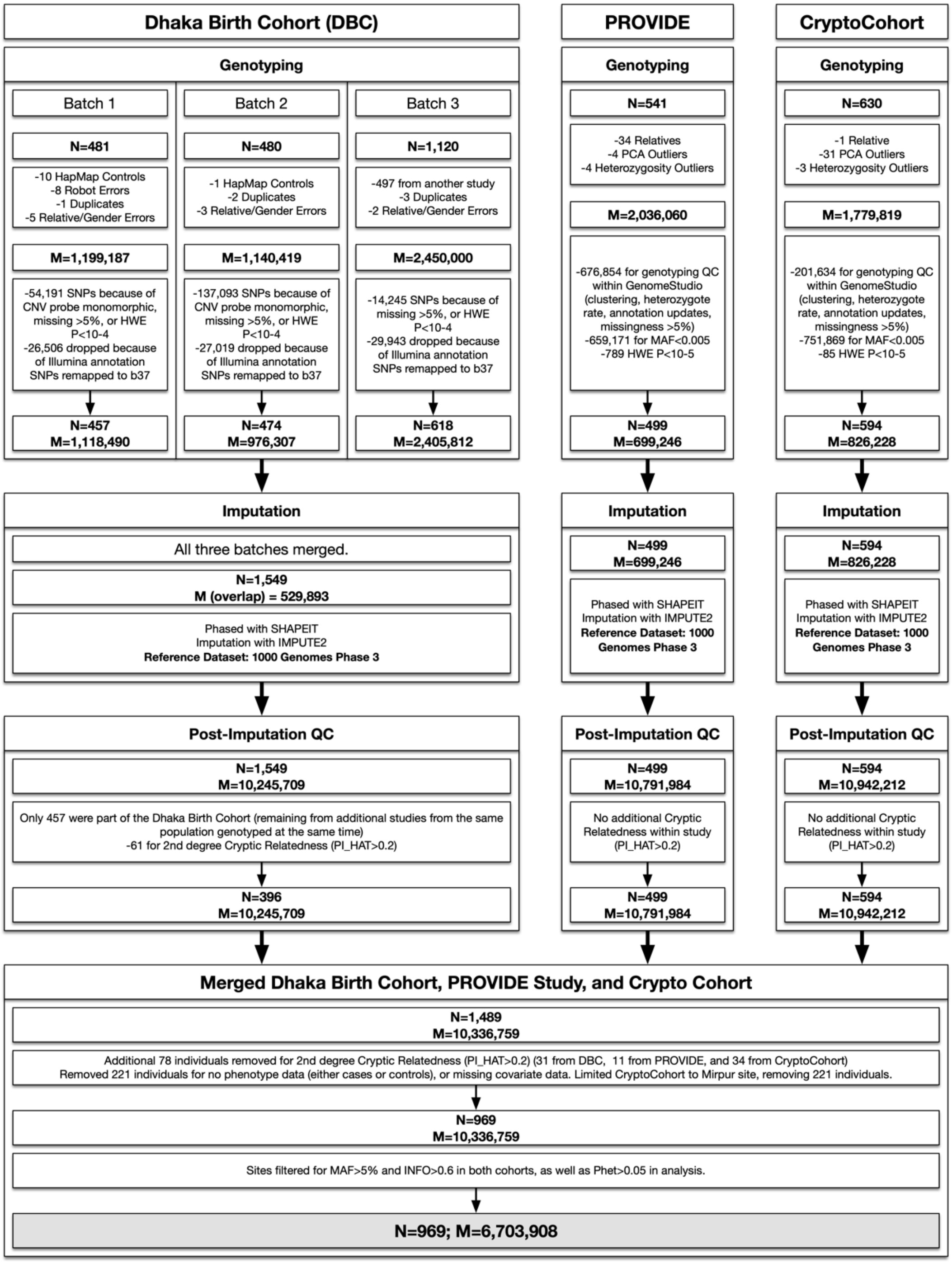


**Supplementary Figure 1: Quality Control workflow for all three cohorts.**

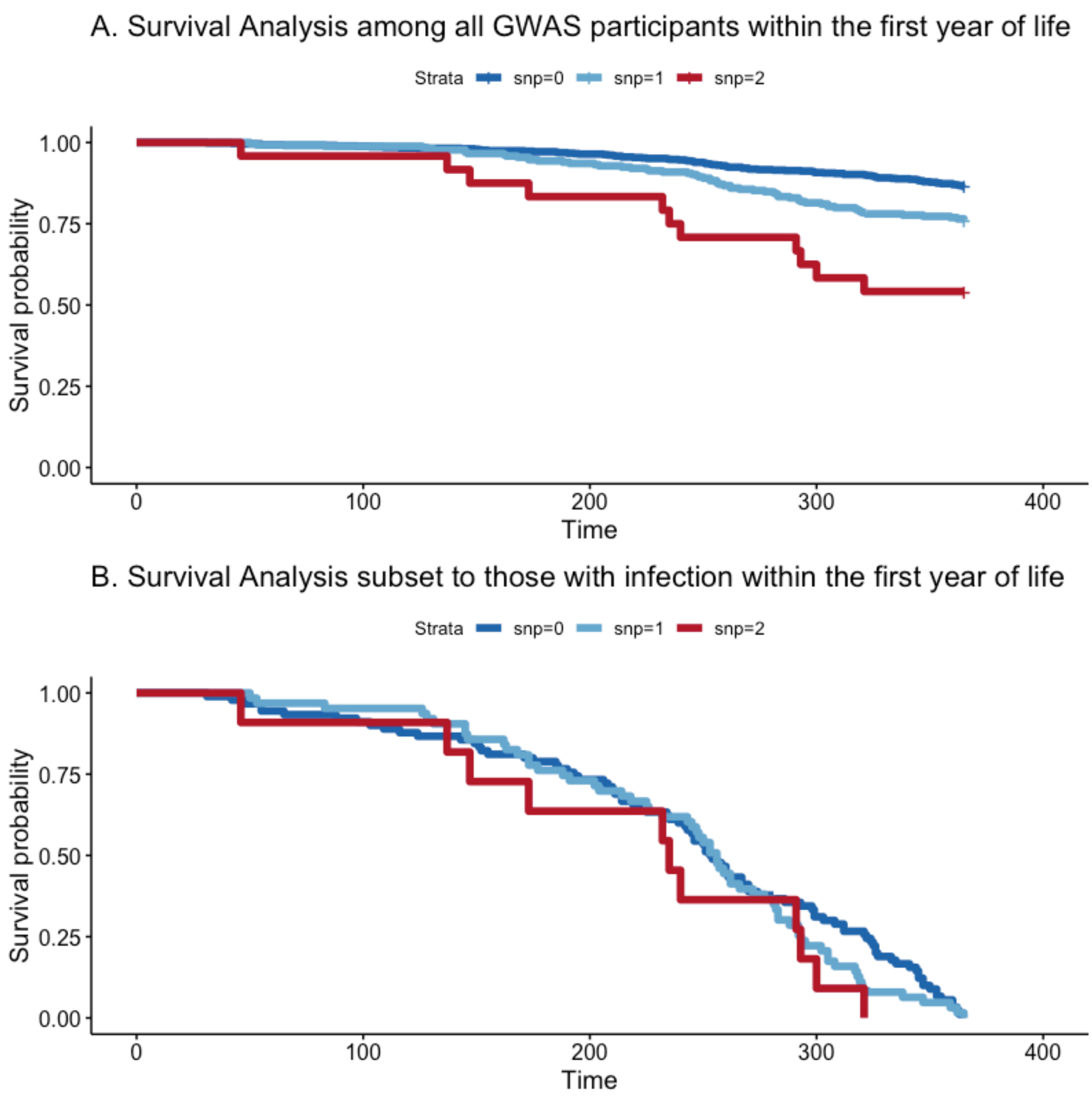


**Supplementary Figure 2: Survival analysis of first episode of cryptosporidiosis by *PRKCA* rs58296998 genotype within the first year of life.**

(A) Survival by genotype within the first year of life (<366 days from birth) among all GWAS participants (both cases and controls). Adjusting for study, we see an additive relationship between the risk allele (T) and earlier infection ($P=6.37x10^{-8}$).

(B) Survival by genotype within the first year of life (<366 days from birth) among only cases in the first year of life. Adjusting for study, we see no additive relationship between the risk allele (T) and earlier infection (P=0.095).

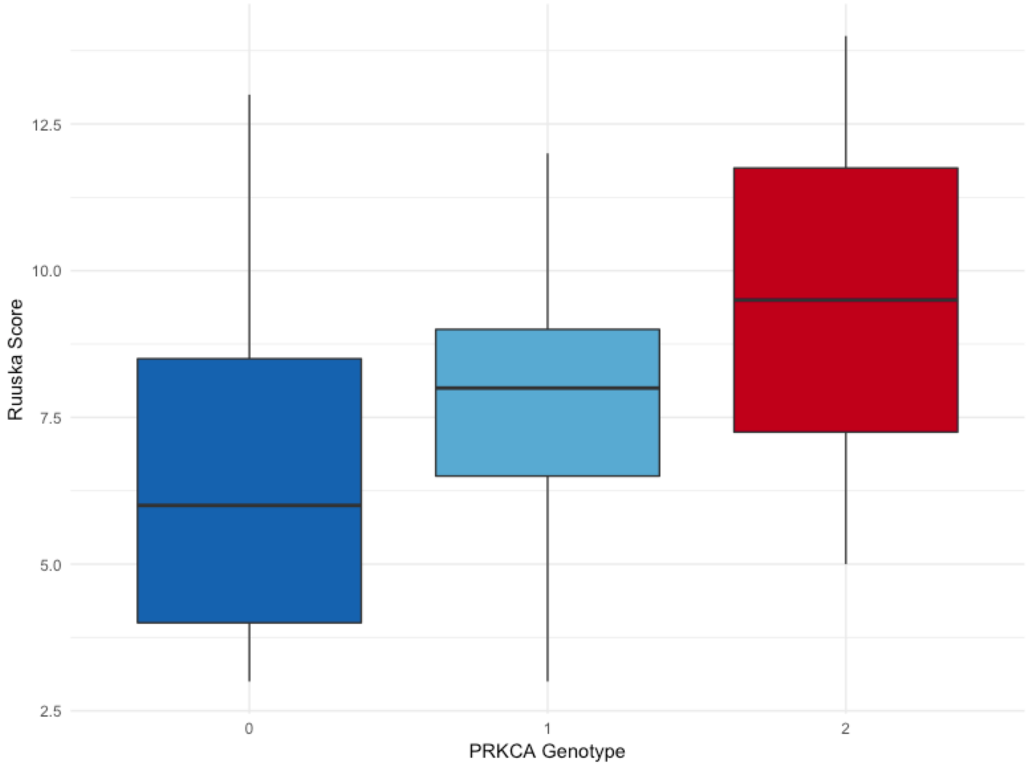


**Supplementary Figure 3: Relationship between genotype for *PRKCA* SNP rs58296998 and severity of diarrhea as determined by Ruuska score within PROVIDE.**

Under an additive model, we see a statistically significant relationship between *PRKCA* genotypes and diarrhea severity (P=0.028).

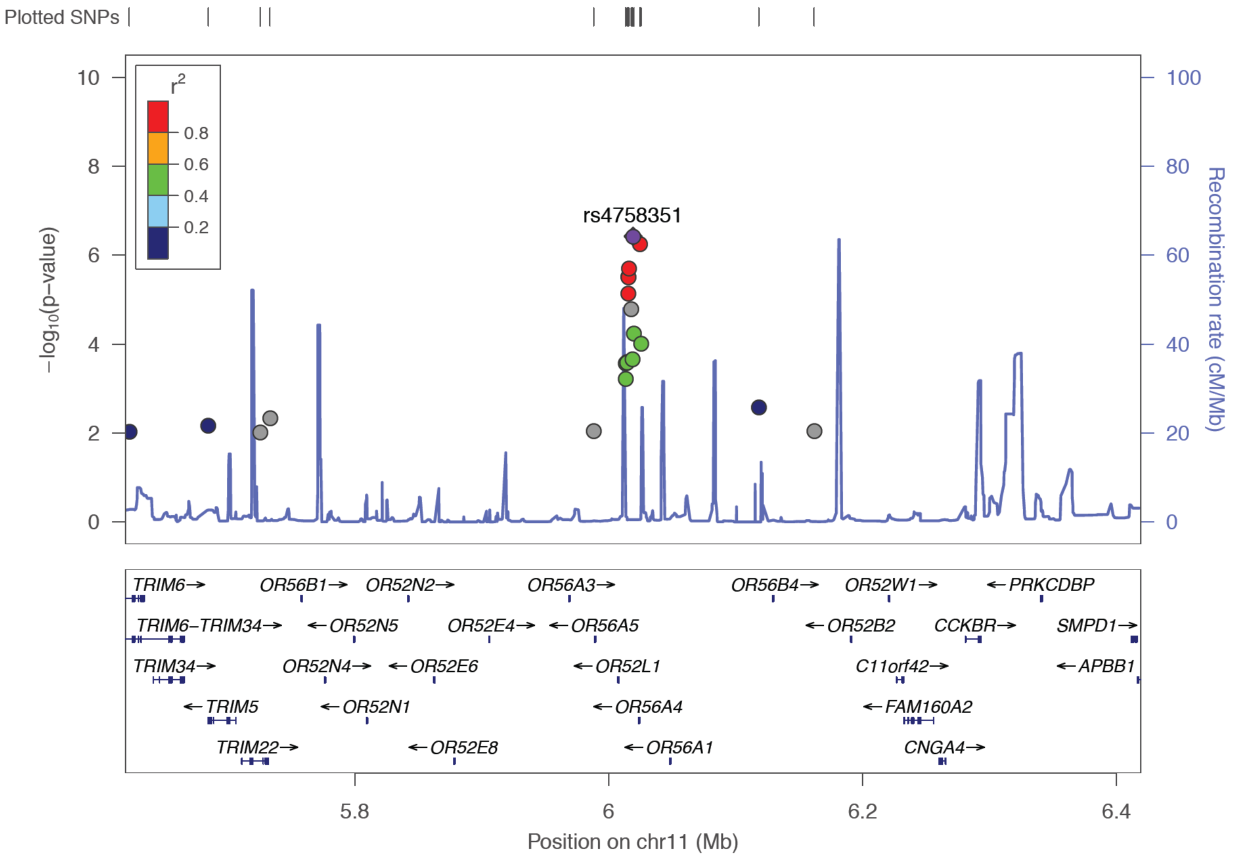


**Supplementary Figure 4: LocusZoom of suggestive signal on chromosome 11 (rs4758351).**

Each dot is a single SNP association from the meta-analysis of the three study-specific logistic regressions adjusting for HAZ(12), the first two study-specific principal components, and batch for DBC. The x-axis is the physical position along chromosome 11 with the gene locations below. The y-axis is the -log P-value from the single SNP association. The fill represents the level of linkage disequilibrium ($r^2$) between the top signal (rs4758351) and surrounding SNPs.

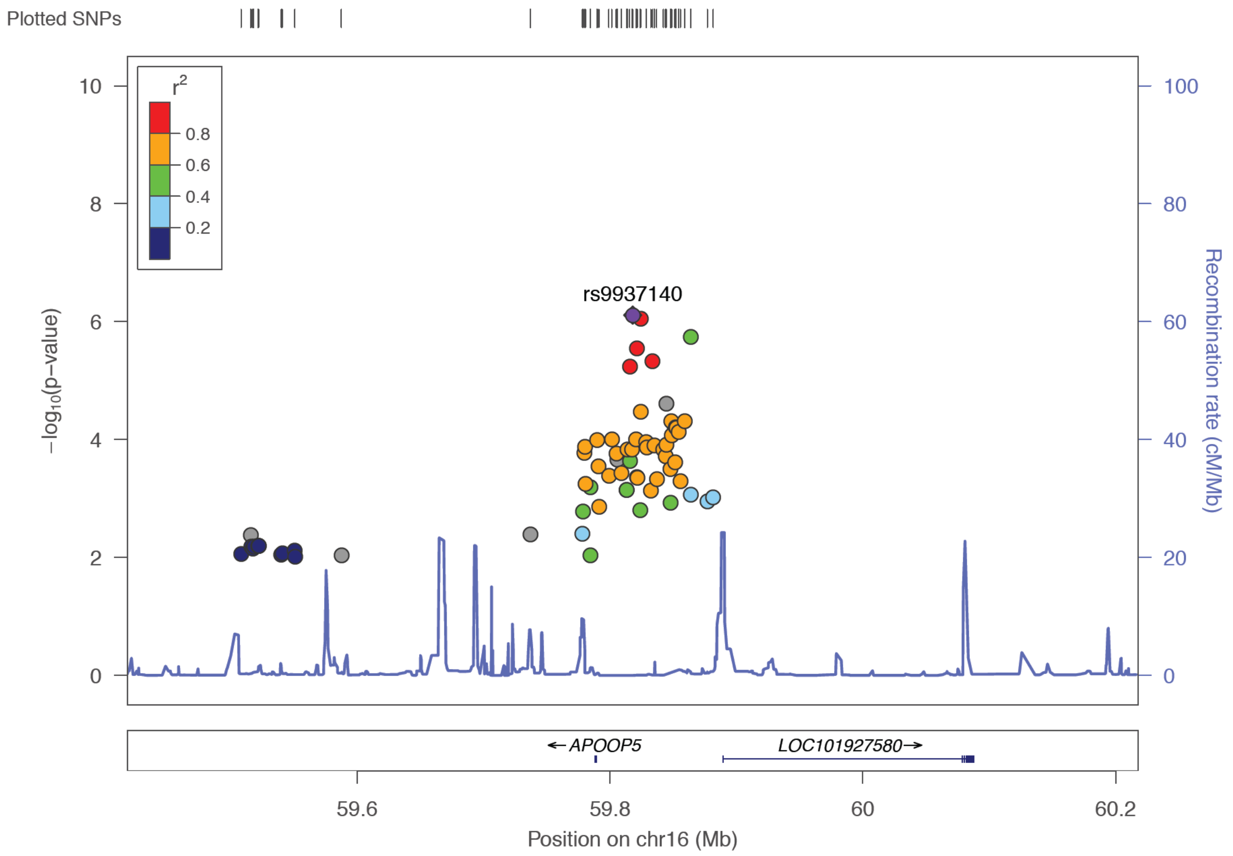


**Supplementary Figure 5: LocusZoom of suggestive signal on chromosome 16 (rs9937140).**

Each dot is a single SNP association from the meta-analysis of the three study-specific logistic regressions adjusting for HAZ(12), the first two study-specific principal components, and batch for DBC. The x-axis is the physical position along chromosome 11 with the gene locations below. The y-axis is the -log P-value from the single SNP association. The fill represents the level of linkage disequilibrium ($r^2$) between the top signal (rs9937140) and surrounding SNPs.

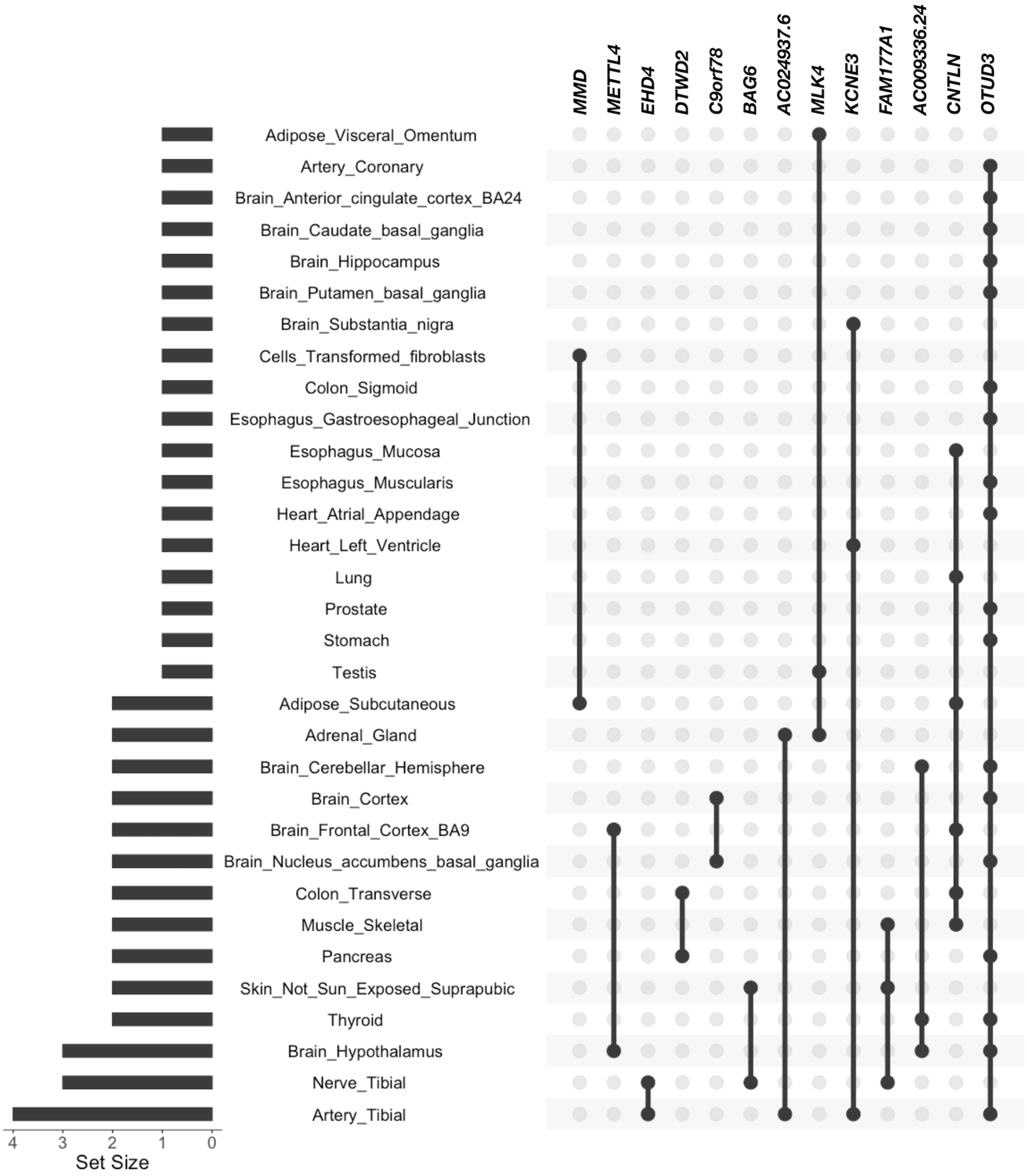


**Supplementary Figure 6: Shared associations for predicted gene expression, filtered for gene-tissue pairs with P<0.001.**

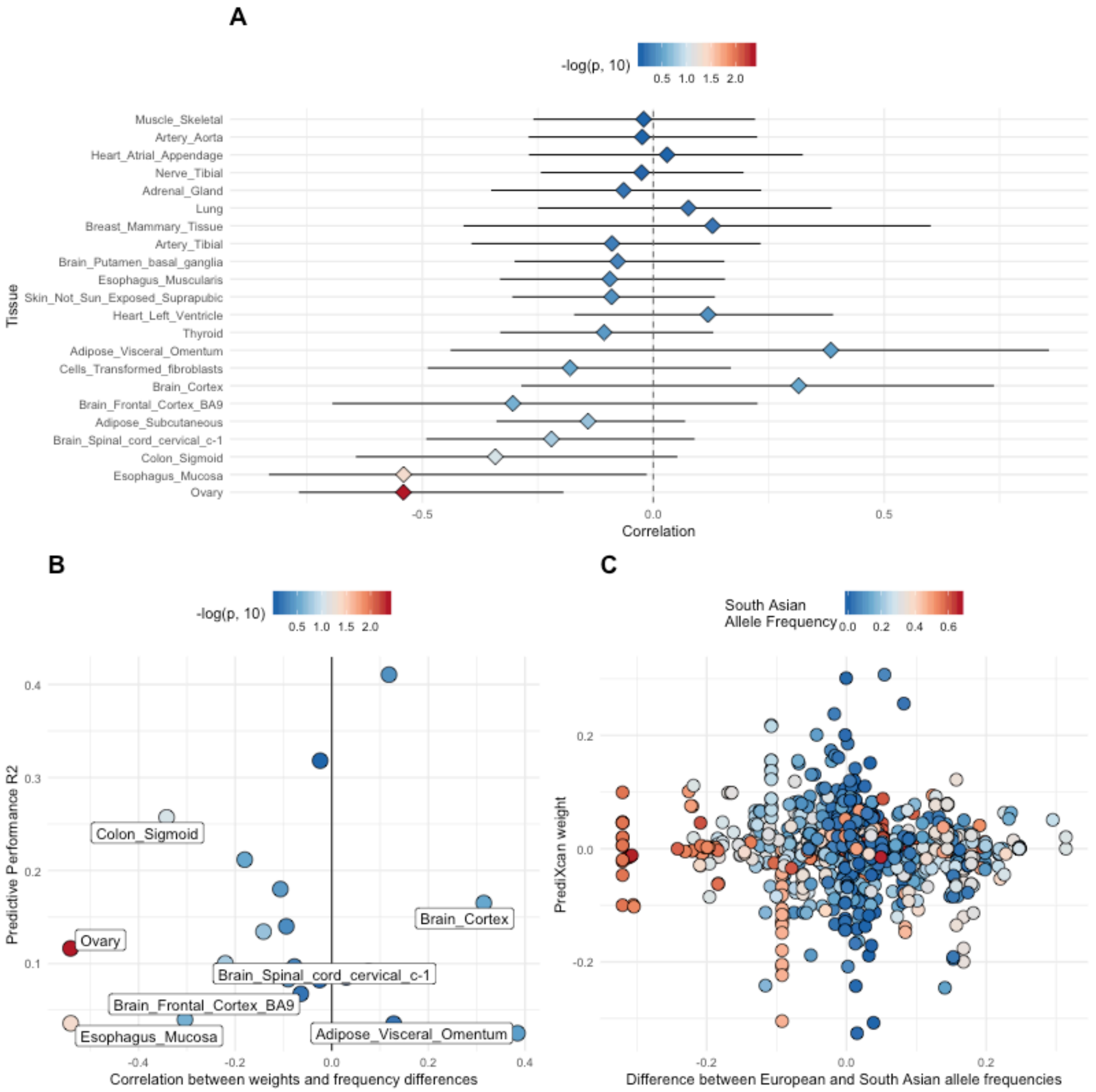


**Supplementary Figure 7: Gene expression prediction characteristics of *PRKCA*.**

(A) Correlation per tissue of differences in allele frequencies between European and South Asian populations with prediXcan weights for *PRKCA*. We see that there is a statistically significant correlation ($P<0.05$) in tissues of interest: colon sigmoid and esophagus mucosa.

(B) Correlation per tissue between weights and frequency differences versus the predictive performance in our participants. The tissues of interest (colon sigmoid, esophagus mucosa) show high correlation and low predictive performance $r^2$. Fill indicates the -Log P-value for correlation.

(C) Difference per SNP between European and South Asian allele frequencies (EUR-SAS) versus the prediXcan weight. Fill indicates the South asian allele frequencies. We note that many of the highest-weighted alleles are low frequency or absent in South Asia.

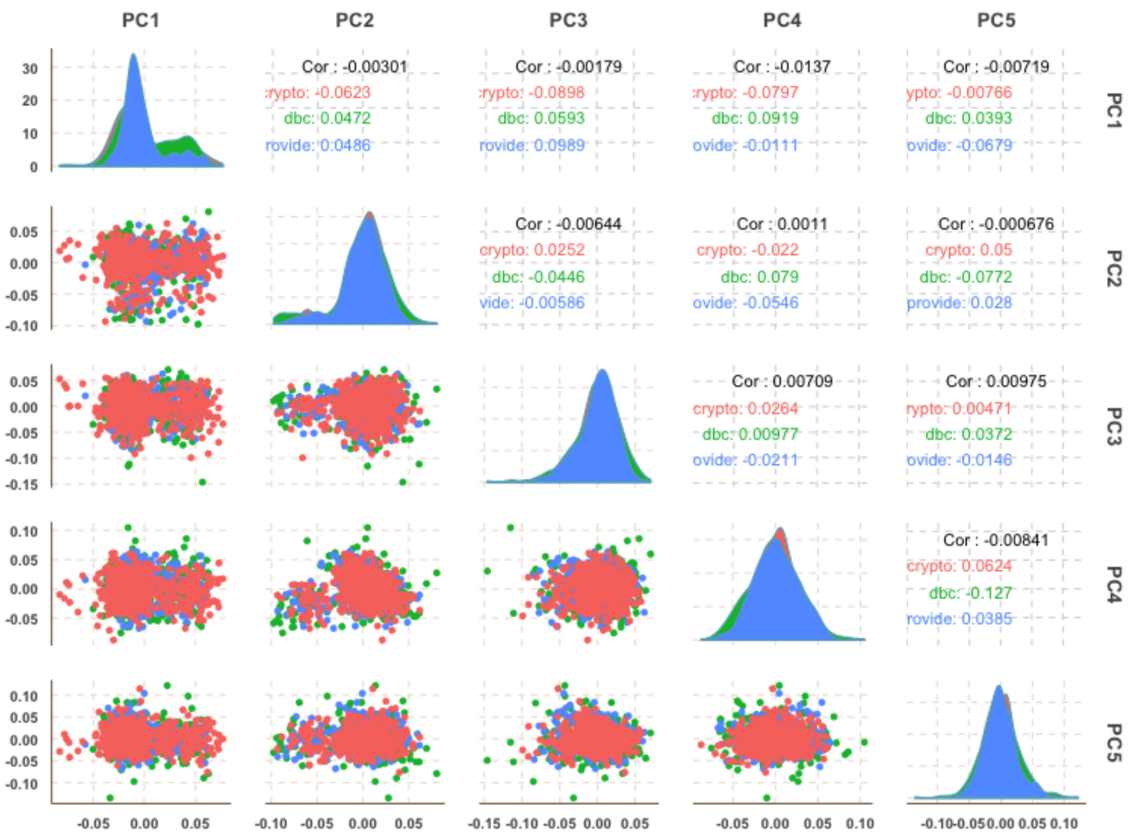


**Supplementary Figure 8: Distribution of three studies for principal components 1-5, colored by study.**
Crypto (red): CryptoCohort, dbc (green): Dhaka Birth Cohort, provide (blue): PROVIDE Study.

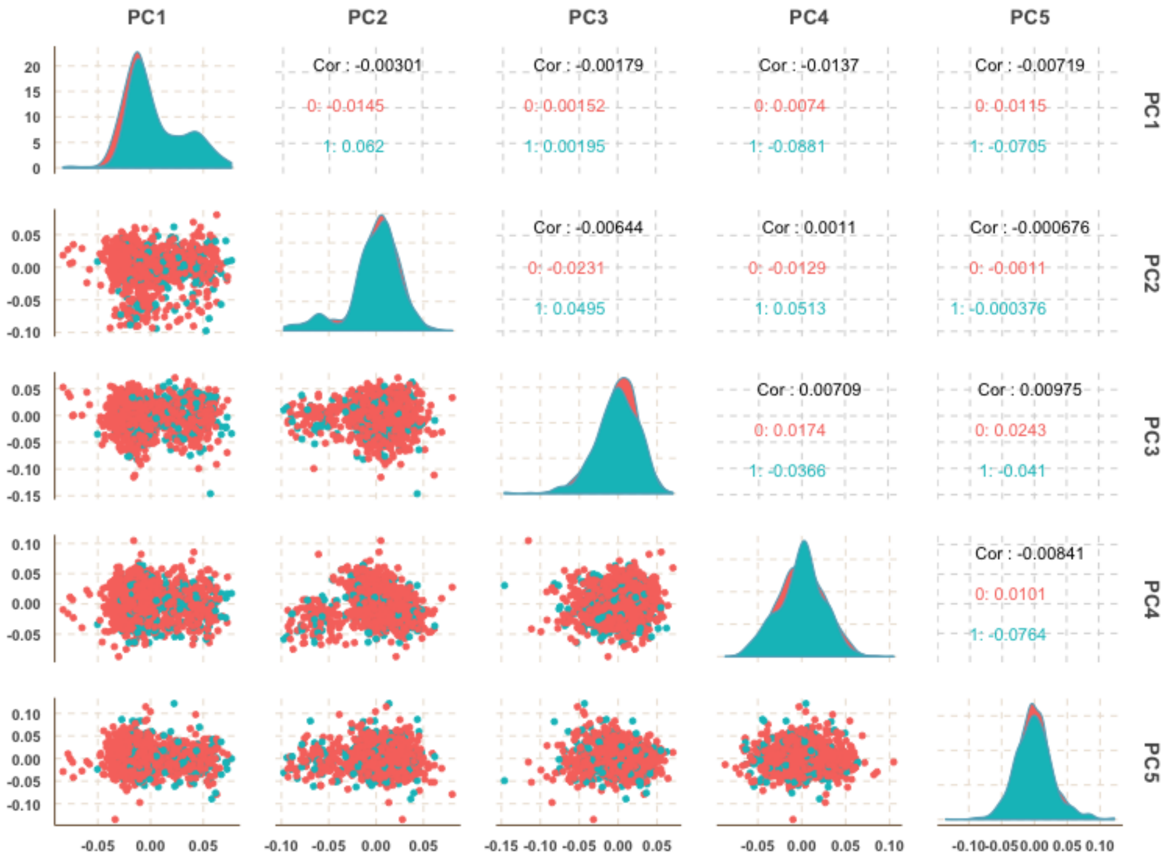


**Supplementary Figure 9: Distribution of three studies for principal components 1-5, colored by case status.**
Cases are shown in blue and controls in red. Only the first principal component was significantly associated with case status.

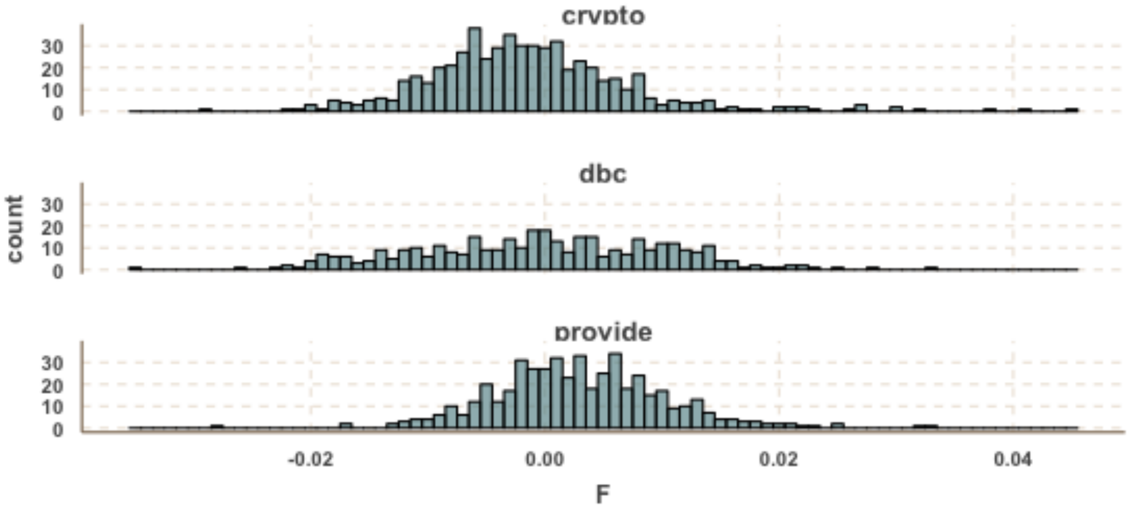


**Supplementary Figure 10: Histogram of heterozygosity distribution by cohort.**
Here we show the distribution of heterozygosity by cohort, crypto: CryptoCohort, dbc: Dhaka Birth Cohort, provide: PROVIDE Study. The x-axis shows F, or the coefficient of heterozygosity.

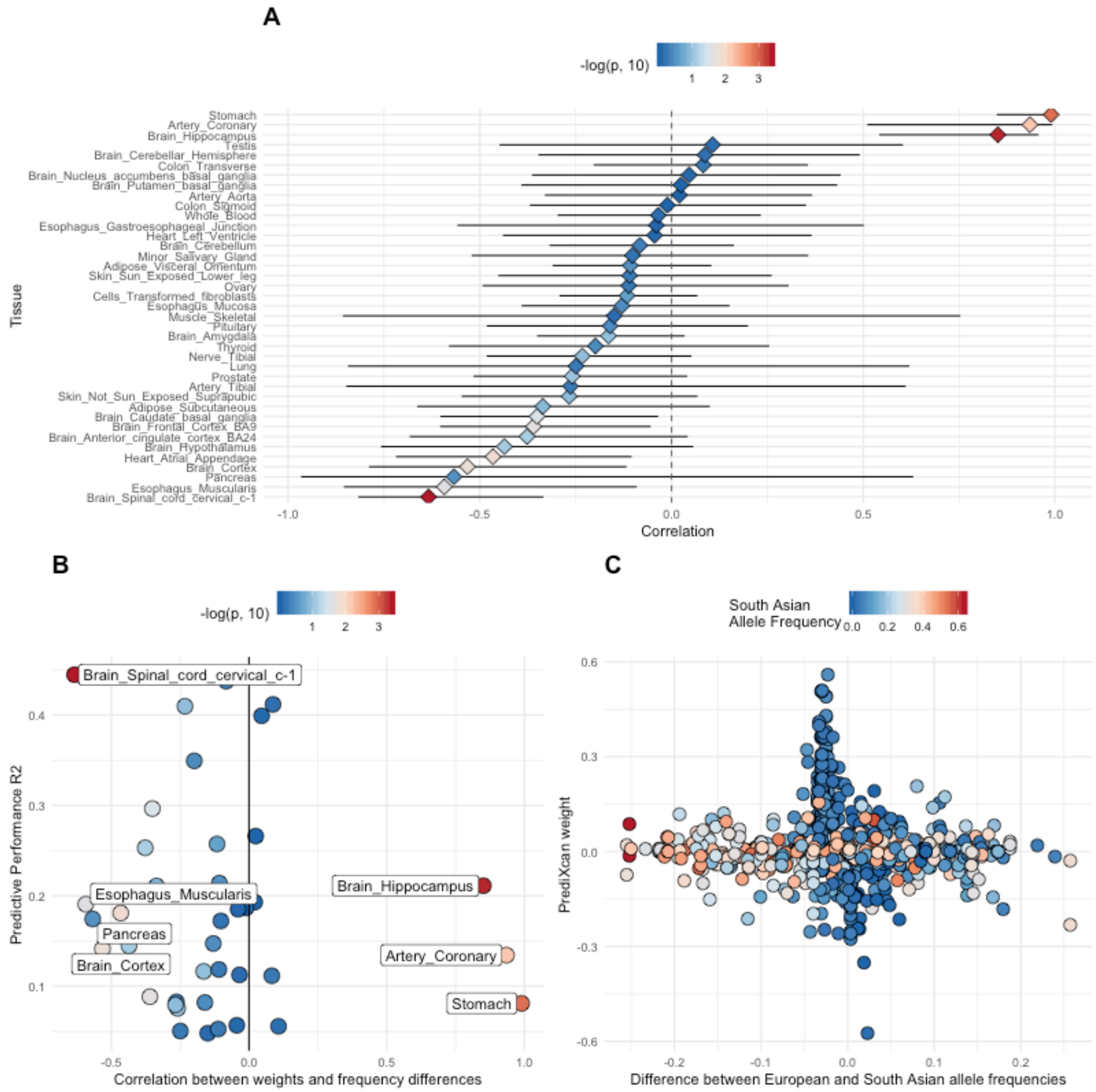


**Supplementary Figure 11: Gene expression prediction characteristics of *OTUD3*.**

(D) Correlation per tissue of differences in allele frequencies between European and South Asian populations with PrediXcan weights for *OTUD3*.

(E) Correlation per tissue between weights and frequency differences versus the predictive performance in our participants. The tissues of interest (colon sigmoid, esophagus mucosa) show high correlation and low predictive performance $r^2$. Fill indicates the -Log P-value for correlation.

(F) Difference per SNP between European and South Asian allele frequencies (EUR-SAS) versus the PrediXcan weight. Fill indicates the South Asian allele frequencies. We note that many of the highest-weighted alleles are low frequency or absent in South Asia.
